# Epigenetic Modulation, Intra-tumoral Microbiome and Immunity in Early Onset Colorectal Cancer

**DOI:** 10.1101/2025.03.28.645992

**Authors:** Ning Jin, Rebecca Hoyd, Ayse Selen Yilmaz, Jiangjiang Zhu, Yunzhou Liu, Malvenderjit Jagjit Singh, Dennis Grencewicz, XiaoKui Mo, Matthew Kalady, Daniel Rosenberg, Caroline E Dravillas, Eric A Singer, John D Carpten, Carlos HF Chan, Michelle L Churchman, Nicholas C Denko, Frances Di Clemente, Rebecca D Dodd, Islam Eljilany, Naomi Fei, Sheetal Hardikar, Alexandra P Ikeguchi, Anjun Ma, Qin Ma, Martin D McCarter, Afaf EG Osman, Gregory Riedlinger, Lary A Robinson, Bryan P Schneider, Ahmad A Tarhini, Gabriel Tinoco, Jane Figueiredo, Yousef Zakharia, Cornelia M Ulrich, Aik Choon Tan, Daniel Spakowicz

## Abstract

**Background:** The incidence of colorectal cancer (CRC) in young adults (age of diagnosis < 50 years old) has been rapidly increasing. Although ∼20% of early-onset (EO) CRC cases are due to germline mutations, the etiology of the majority of EOCRC cases remains poorly understood. Non-genetic factors such as environmental exposure and lifestyle changes are likely to have a direct link to the increased incidence of sporadic EOCRC. We hypothesize that such factors may be observable as differences in the EOCRC epigenome, microbiome and immunome. We sought to address this by comparing differences in DNA methylation from the cohort of colorectal cancer patients in The Cancer Genome Atlas (TCGA). Further, we carefully identified intra-tumoral microbes from TCGA and two other datasets and then related the microbes to EOCRC status and deconvolved immune cell abundances. We found that DNA methylation (DNAm) age acceleration by12 years when compared with average-onset CRC (AOCRC) patients. Differentially methylated sites associated with genes are related to CREB signaling in neurons, G protein coupled receptor signaling, phagosome formation and S100 family signaling. These differences were validated in the gene expression from TCGA and a second, larger real-world dataset from the Oncology Research Information Exchange Network (ORIEN). However, no consistent differences were observed in the intra-tumor microbes between EOCRC and AOCRC. Interestingly, the most abundant microbes interacted with the immune systems differently between the EOCRC and AOCRC tumors, characterized by more, larger, positive correlations in EOCRC. These data suggest epigenetic modulation and accelerated aging may play a key role in the development of EOCRC.

**SIGNIFICANCE:** We investigated whether environmentally driven factors contribute to early-onset colorectal cancer (EOCRC). We observed accelerated epigenetic aging in EOCRC and epigenetic changes associated with chronic inflammation. Tumor immune cell abundances correlated more strongly with microbes in EOCRC than average-onset CRC. These data suggest a dysregulation of immune response in EOCRC, driving chronic inflammation and tissue aging.

## INTRODUCTION

Colorectal cancer (CRC) is the third most common cause of cancer deaths both in men and women in the United States (US). Although CRC incidence and mortality have decreased steadily among patients older than 65 years, the incidence of CRC in younger adults (< 50 years old) has been rapidly increasing by 2% per year since the early 1990s.^1^ Approximately 20% of early-onset CRC (EOCRC) cases are attributed to germline DNA mutations that are related to familial cancer syndromes.^2–4^ The surge in CRC incidence in young adults is particularly alarming as the overall CRC frequency has decreased. There is no consensus on whether EOCRC is characterized by distinct molecular features compared to average-onset CRC (AOCRC), defined as patients with CRC diagnosis at 50 years old or later. EOCRC is characterized by the advanced stage at diagnosis, poor cell differentiation, higher prevalence of signet ring cell histology, and distal location of the primary tumor (especially the rectum).^5, 6^ Available data suggest that survival is worst in patients younger than 30 years old, and sporadic CRC among the youngest patients represents a significant clinical challenge.^7^

The etiology of sporadic EOCRC is poorly understood. Epidemiology studies estimate that approximately 50–60% of CRC cases in the US are attributable to modifiable risk factors, such as smoking, alcohol, obesity, physical inactivity, and a Western diet.^8–12^ In one study, African Americans were found to have a higher CRC risk compared with South Africans living in rural areas.^13^ The higher CRC rates in African Americans were associated with diets with higher animal protein and fat, and lower fiber consumption.^13^ Diet also has a major impact on the composition and function of gut microbiota, which decompose and ferment dietary fibers to produce microbial metabolites, such as short-chain fatty acids (SCFAs), polyphenols, or vitamins.

Changes in the gut microbiome composition (including bacteria, viruses, and fungi) may lead to various diseases, including irritable bowel syndrome, Crohn’s disease, and CRC.^14–18^ A diet with high fat and low fiber has been associated with disruption of intestinal bacterial homeostasis, which is termed intestinal dysbiosis.^8, 12, 19^ Studies have found that intestinal microbiome composition is more diverse in healthy individuals than in CRC patients. Additionally, specific microbes have been linked to different stages of colorectal neoplasia, from adenomatous polyps to early-stage cancer to metastatic disease.^20–25^ For example, CRC-associated taxa such as *Fusobacterium* (anaerobic gram-negative bacteria) were frequently found in CRC tumors compared with adjacent normal colon tissue.^26^ Animal studies showed *Fusobacterium* can activate β-catenin signaling, promoting oncogenesis.^27^ Other pathogenic microbes, such as enterotoxigenic *Bacteroides fragilis* (ETBF), can cause T helper 17 (Th17)-driven inflammation, leading to colonic tumor development.^28^ Therefore, changes in the intestinal microbiome and microbial metabolites may facilitate environmental risk factors to initiate and promote CRC.^29, 30^

Numerous studies have found that microbial metabolites play important roles in modulating the host immune system.^31–34^ Intestinal microbiota are essential in developing and regulating innate and adaptive immunity.^35, 36^ Local immunity is promoted by recognizing pathogen-associated molecular patterns (PAMPs) by pattern-recognition receptors (PRRs) in intestinal epithelial cells and innate immune cells within the gut. Bacterial and microbial metabolites can activate dendritic cells (DCs), which migrate to the draining lymph nodes to activate naïve T cells to form either T regulatory (Treg) or Th17 cells.^37^ To maintain intestinal homeostasis, the immune system must tolerate antigens derived from commensal microbiota. Tolerance is achieved in part via the actions of Tregs, which produce the anti-inflammatory cytokine interleukin-10 (IL-10). In the setting of acute infection due to opportunistic pathogenic bacteria, the delicate balance of the commensal bacteria is disrupted, ^38^ leading to the breakdown of the mucosal barrier and the translocation of gut bacteria to local lymph nodes and the peripheral circulation. Consequently, Th17 cells produce the pro-inflammatory cytokine IL-17 and recruit neutrophils from the bloodstream, generating a profound inflammatory state.^39^ The chronic inflammatory state leads to the activation of key pro-survival and pro-proliferative signaling pathways via NF-κB, resulting in aberrant proliferation, epithelial cell transformation, and ultimately the development of CRC.^40, 41^ Therefore, the unique immune response to specific or pathogenic microbes may contribute to the development of EOCRC.

In addition to promoting CRC, microbial metabolites may influence the host epigenome, either by regulating the activity of epigenetic enzymes or by altering the abundance of cofactors needed for epigenetic modifications.^42, 43^ For instance, butyrate, one of the SCFAs produced by microbes, may inhibit histone deacetylases and colonic cell proliferation.^44^ Folic acid and other B vitamins from green leafy vegetables are the essential methyl donors required to provide substrates for DNA methylation. Decreased dietary folate intake may deplete the levels of S-adenosyl methionine (SAM), resulting in DNA hypomethylation, which may lead to the activation of proto-oncogenes and chromosome instability.^42, 43^ regulates gene transcription by either gene activation or silencing through DNA methyltransferases (DNMTs), which converts cytosine to 5-methylcytosine.^45^ DNA methylation at CpG islands in promoters are generally repressive markers that prevent gene transcriptional activity by impeding recruitment of transcriptional machinery. On the other hand, DNA methylation within intronic regions may cause increased transcription in a tissue-specific manner.^46^ Hypermethylation of specific regions, such as the CpG islands of tumor suppressor genes, plays an important role in carcinogenesis for many types of cancers, including CRC.^43, 47–49^ Studies have suggested that methylation changes may be the earliest alterations in the polyp-to-cancer transformation, suggesting that methylation may play a key role in CRC tumorigenesis.^50–53^ In fact, a study compared 118 EOCRC cases with 225 AOCRC cases and found hypomethylation in long interspersed nuclear element-1 (LINE-1, the repetitive genomic elements) in EOCRC.^54^ However, few studies have investigated the genome-wide epigenetic alterations that may underlie EOCRC.

In this study, our goal was to obtain further insight into the mechanisms associated with EOCRC. We hypothesized that EOCRC may be reflected by alterations in the intra-tumoral microbiome and DNA methylation, which are the markers for environmental changes. We aimed to characterize the methylation signatures and investigate the unique distribution of the intra-tumoral microbiome and interactions between the microbiome and tumor-infiltrating lymphocytes (TILs) in EOCRC.

## METHODS

### TCGA Dataset

A total of 358 CRC cases, including 54 cases of EOCRC (age at diagnosis < 50 years) and 304 cases of AOCRC (age at diagnosis ≥ 50 years), with matched methylation array (Infinium HM450), and RNA sequencing (RNA-seq) (Illumina HiSeq) data from colon adenocarcinomas (COAD) and rectal adenocarcinomas (READ), and clinicopathological information of each patient, were extracted from The Cancer Genome Atlas (TCGA). The TCGA biolinks BioConductor package was used to download and analyze both cohorts.^55^ We used the TCGA biolinks BioConductor package to download Infinium HM450 data and analyzed EOCRC and AOCRC cohorts.^56^ After normalization and adjustment of the batch effect with the ComBat package, the average β value of methylated CpG sites for each patient was calculated. We applied the batch effect removal tool ComBat to the level-3 methylation beta values.^57^ We performed a differential methylated CpG analysis to estimate the difference in DNA methylation for the probes between groups and their significance value using the Wilcoxon test and the Benjamini-Hochberg adjustment method. An adjusted *P* < 0.05 was considered to be a differentially methylated CpGs.

### ORIEN Dataset

The Oncology Research Information Exchange Network (ORIEN) is an alliance of 18 cancer centers in the US established in 2014. All ORIEN alliance members utilize a standard Total Cancer Care (TCC) protocol. As part of the TCC study, participants agree to have their clinical data followed over time, undergo germline and tumor sequencing, and be contacted by their provider if an appropriate clinical trial or other study becomes available. TCC is a prospective cohort study with a subset of patients enrolled in the ORIEN Avatar program, including research use only (RUO)-grade whole-exome tumor sequencing, RNA sequencing, germline sequencing, and deep longitudinal clinical data collection with lifetime follow-up.

Nationally, over 325,000 participants have enrolled in TCC. Aster Insights, ORIEN’s commercial and operational partner, harmonizes all abstracted clinical data elements and molecular sequencing files into a standardized, structured format to enable aggregation of deidentified data for sharing across the network. The Ohio State University (OSU) Institutional Review Board (IRB) approved the study protocol (2015H0088), which is registered at ClinicalTrials.gov (NCT02482610).

ORIEN Avatar specimens undergo nucleic acid extraction and sequencing at HudsonAlpha (Huntsville, AL) or Fulgent Genetics (Temple City, CA). For frozen tissue DNA extraction, Qiagen QIASymphony DNA purification is performed, generating a 213-bp average insert size. For frozen and optimal cutting temperature (OCT) tissue RNA extraction, Qiagen RNAeasy plus mini kit is performed, generating a 216-bp average insert size. For formalin-fixed paraffin-embedded (FFPE) tissue, a Covaris Ultrasonication FFPE DNA/RNA kit is utilized to extract DNA and RNA, generating a 165-bp average insert size. RNA-seq is performed using the Illumina TruSeq RNA Exome with single library hybridization, cDNA synthesis, library preparation, and sequencing (100-bp paired reads at Hudson Alpha, 150-bp paired reads at Fulgent) to a coverage of 100 million total reads/50 million paired reads.

Our study included 453 ORIEN Avatar patients with colon or rectal cancer who were consented to the TCC protocol from the participating member sites of ORIEN. We filtered out all cases with microsatellite instability status in both TCGA and ORIEN datasets.

### Data Processing for RNA Sequencing

RNA-seq tumor pipeline analysis is processed according to the following workflow using GRCh38/hg38 human genome reference sequencing and GenCode build version 32. Adapter sequences are trimmed from the raw tumor sequencing FASTQ file. Adapter trimming via k-mer matching is performed along with quality trimming and filtering, contaminant filtering, sequence masking, guanosine/cytosine filtering, length filtering, and entropy filtering. The trimmed FASTQ file is used as input to the read alignment process. The tumor adapter-trimmed FASTQ file is aligned to the human genome reference (GRCh38/hg38) and the Gencode genome annotation v32 using the STAR aligner. The STAR aligner generates multiple output files for gene fusion prediction and gene expression analysis. RNA expression values are calculated and reported using estimated mapped reads, fragments per kilobase of transcript per million mapped reads (FPKM), and transcripts per million mapped reads (TPM) at both the transcript and gene levels based on transcriptome alignment generated by STAR. The RSEM pipeline and gene expressions (GEs)were quantified as TPM. GEs were log2(TPM+1) transformed, and downstream analyses were performed using the GE matrix.

For gene expression analysis, primary alignment is performed against the human genome reference GRCh38. Gene expression values were quantified using the featureCounts tool of the Subread package v1.5.1 in unstranded mode for genes described by the GENCODE annotation. Differential expression analysis was performed using Limma/voom.^110^ The functional analyses were generated using QIAGEN Ingenuity pathway analysis (IPA) (QIAGEN Inc., https://digitalinsights.qiagen.com/IPA).^58^ Cutoff values for the core analysis in IPA include *P*-value < 0.05 and |Fold change| > 1.5.

### Microbe Alignment and Quantification with {exotic}^59^

We utilized the tool {exotic},^59^ which takes raw RNA-seq data (in this case, from the ORIEN and TCGA datasets) and carefully aligns it to both human and non-human reference genomes to identify low-abundance microbes. Models of association were analyzed based on each of the subgroups as well as all of the samples (the “all” group). We performed Cox proportional hazards regression to identify the microbes associated with overall survival (OS). We evaluated 453 RNA-seq samples from ORIEN and then validated those associations on an independent dataset of 358 samples from TCGA as previously described.^59^

### Methylation Array and Epigenetic Clock Analysis in TCGA

DNA methylation profiles were downloaded from the Illumina Infinium Human Methylation 450 platform in TCGA. DNA methylation values are reported as beta values for each CpG probe in each sample using Illumina BeadStudio software. Beta values are continuous variables between 0 and 1, representing the ratio of the intensity where higher beta values suggest a higher level of DNA methylation (hypermethylation) and lower beta values suggest a lower level of DNA methylation (hypomethylation). All the methylation values are batch corrected using the Combat function from the sva package.^108^ Then, the difference of mean methylation between EOCRC and AOCRC groups for each probe are identified using the TCGAanalyzeDMR function in the TCGABiolinks package.^109^ The *P* value is calculated using the Wilcoxon test and adjusted using the Benjamini-Hochberg method. The differentially methylated sites are identified using a difference in methylation value |Δ beta| > 0.1 and a false discovery rate (FDR) adjusted *P* value of <0.05.

We calculated the epigenetic ages of TCGA EOCRC and AOCRC cohorts. The specialized R library Enmix^60^ is used to generate the three well-established epigenetic scores: Horvath,^61^ Hannum,^62^ and PhenoAge.^63^ We calculated the Δ age difference between each epigenetic age and the chronological age for both AOCRC and EOCRC cohorts.

### Deconvolution of RNA Sequencing to Characterize TILs

Reads that aligned to the human reference genome were used to estimate gene transcript abundances. The count table was deconvolved to the abundances of immune cell types. TILs were fit to the gene expression set using a support vector regression method that performs variable selection on the signature gene set to prevent overfitting. Reference cell-type signatures of individual lymphocytes were aggregated.

### Data Availability

The OSU IRB approved data access in an Honest Broker protocol (2015H0185) and TCC protocol (2013H0199) in coordination with Aster Insights. The processed data generated in this study are publicly available in Gene Expression Omnibus through the BioProject PRJNA856973. Analysis scripts and data to regenerate all figures and tables are available at: https://github.com/spakowiczlab/exorien-exogieo.

## RESULTS

### EOCRC Patient Characteristics from Two Datasets

We characterized the methylation, gene expression patterns, and microbiome signature in EOCRC for both TCGA and ORIEN datasets. We define EOCRC as age at diagnosis < 50 and AOCRC as age at diagnosis ≥ 50. We collected an Infinium HM450 methylation array with matched RNA-seq data from TCGA, comprising a total of 358 CRC cases, including 54 EOCRC and 304 AOCRC cases (**Table 1**).

**Table 1:** Patient Characteristics with COAD and READ tumors in TCGA, stratified by the age of onset.

|  | EO | AO | p |
| --- | --- | --- | --- |
| **N** | **54** | **303** |  |
| **Sex = Male (%)** | **28 (51.9)** | **167 (55.1)** | **0.768** |
| **Race (%)** |  |  | **0.246** |
| American Indian or Alaska Native | 0 (0.0) | 1 (0.3) |  |
| Asian | 3 (5.6) | 8 (2.6) |  |
| Black or African American | 11 (20.4) | 40 (13.2) |  |
| Not reported | 1 (1.9) | 23 (7.6) |  |
| White | 39 (72.2) | 231 (76.2) |  |
| **Primary Diagnosis Site (%)** |  |  | **0.454** |
| Colon | 38 (70.4) | 227 (74.9) |  |
| Connective, or soft tissues | 0 (0.0) | 2 (0.7) |  |
| Rectosigmoid Junction | 10 (18.5) | 34 (11.2) |  |
| Rectum | 6 (11.1) | 40 (13.2) |  |
| **Tumor Location (%)** |  |  | **0.343** |
| Left | 30 (55.6) | 140 (46.2) |  |
| Other | 8 (14.8) | 42 (13.9) |  |
| Right | 16 (29.6) | 121 (39.9) |  |
| **Tumor Stage (%)** |  |  | **0.046** |
| Stage I | 6 (11.5) | 46 (16.1) |  |
| Stage II | 13 (25.0) | 114 (39.9) |  |
| Stage III | 20 (38.5) | 88 (30.8) |  |
| Stage IV | 13 (25.0) | 38 (13.3) |  |
| **Histology (%)** |  |  | **0.238** |
| Adenocarcinoma with mixed subtypes | 1 (1.9) | 1 (0.3) |  |
| Adenocarcinoma, NOS | 42 (77.8) | 256 (84.5) |  |
| Mucinous adenocarcinoma | 10 (18.5) | 32 (10.6) |  |
| Papillary adenocarcinoma, NOS | 0 (0.0) | 2 (0.7) |  |
| Tubular adenocarcinoma | 1 (1.9) | 12 (4.0) |  |

Then, we obtained 453 CRC cases with RNA-seq from ORIEN, including 120 EOCRC and 333 AOCRC cases (**Table 2**). In both TCGA and ORIEN datasets, EOCRC patients differed from AOCRC patients by cancer stage, with EOCRC patients having a more advanced stage at initial diagnosis (*P*-value = 0.046 from TCGA; *P*-value = 0.018 from ORIEN). In ORIEN datasets, EOCRC patients were found to have more left-sided primary tumor locations than right-sided tumor locations (*P*-value = 0.008). Histological subtypes, race, and sex were not significantly different between EOCRC and AOCRC patients. Our observance of EOCRC presenting with advanced cancer stage and left-sidedness in our datasets is consistent with other research groups.^5, 6^

**Table 2:**
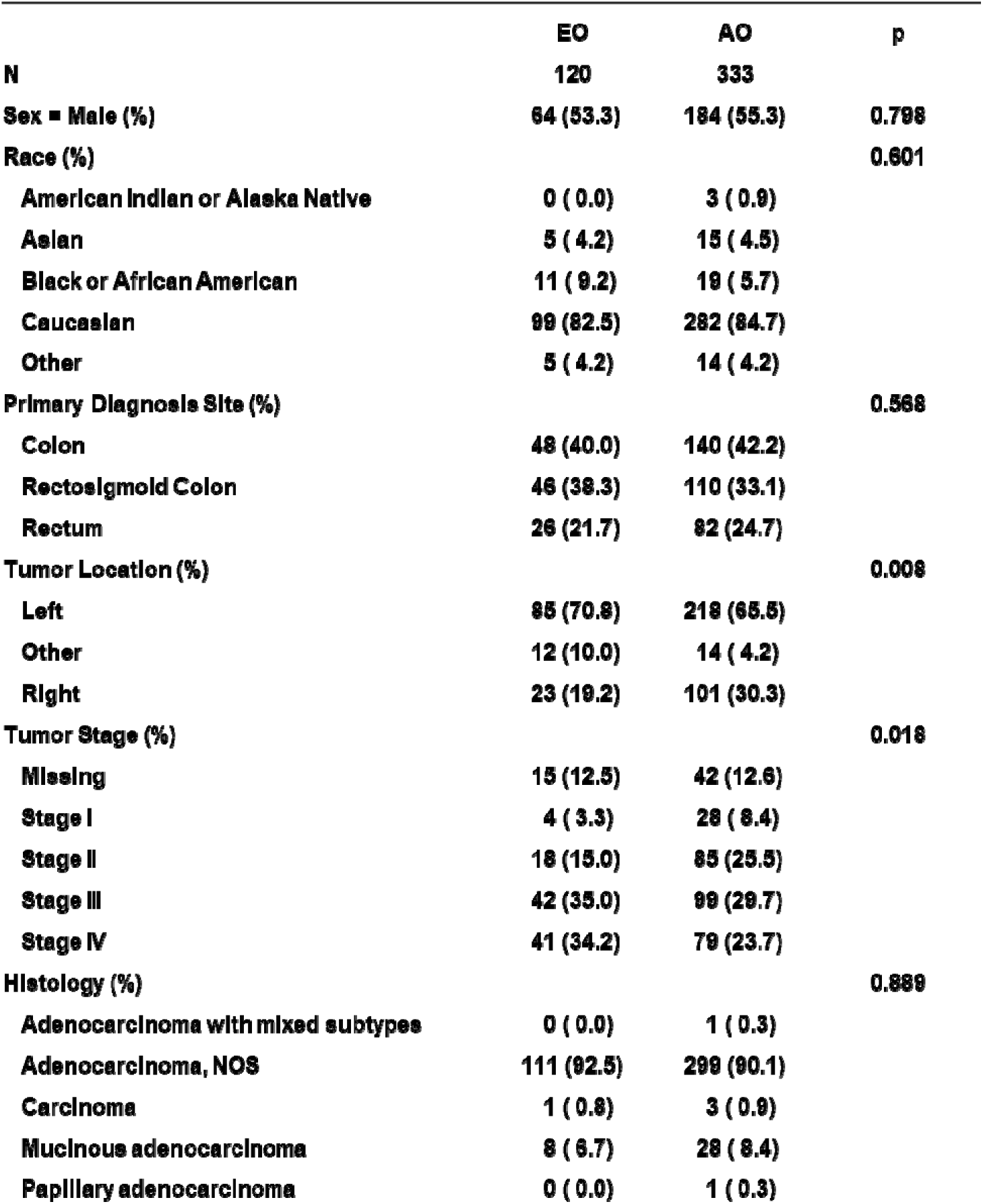
Patient Characteristics with CRC tumors in TCC, stratified by the age of onset.

### Epigenetic Signatures and Accelerated Aging in EOCRC

We analyzed differential methylated sites (DMSs) in the EOCRC and AOCRC groups using an Infinium HM450 methylation array in TCGA. A DNA methylation heatmap demonstrated the methylation patterns in EOCRC and AOCRC with the individual cases (patients) **(Fig. 1A)**. The volcano plot indicated more hypomethylated DMSs in the EOCRC group **(Fig. 1B**, per site differences and *P*-values in **Supplementary Table S1)**. However, the mean global methylations were not significantly different between the two groups (EOCRC and AOCRC patients were 0.46 ± 0.028 and 0.47 ± 0.027, *P*-value=0.13, 2-sample *t*-test) (**Figure 1C**).

**Figure 1.**
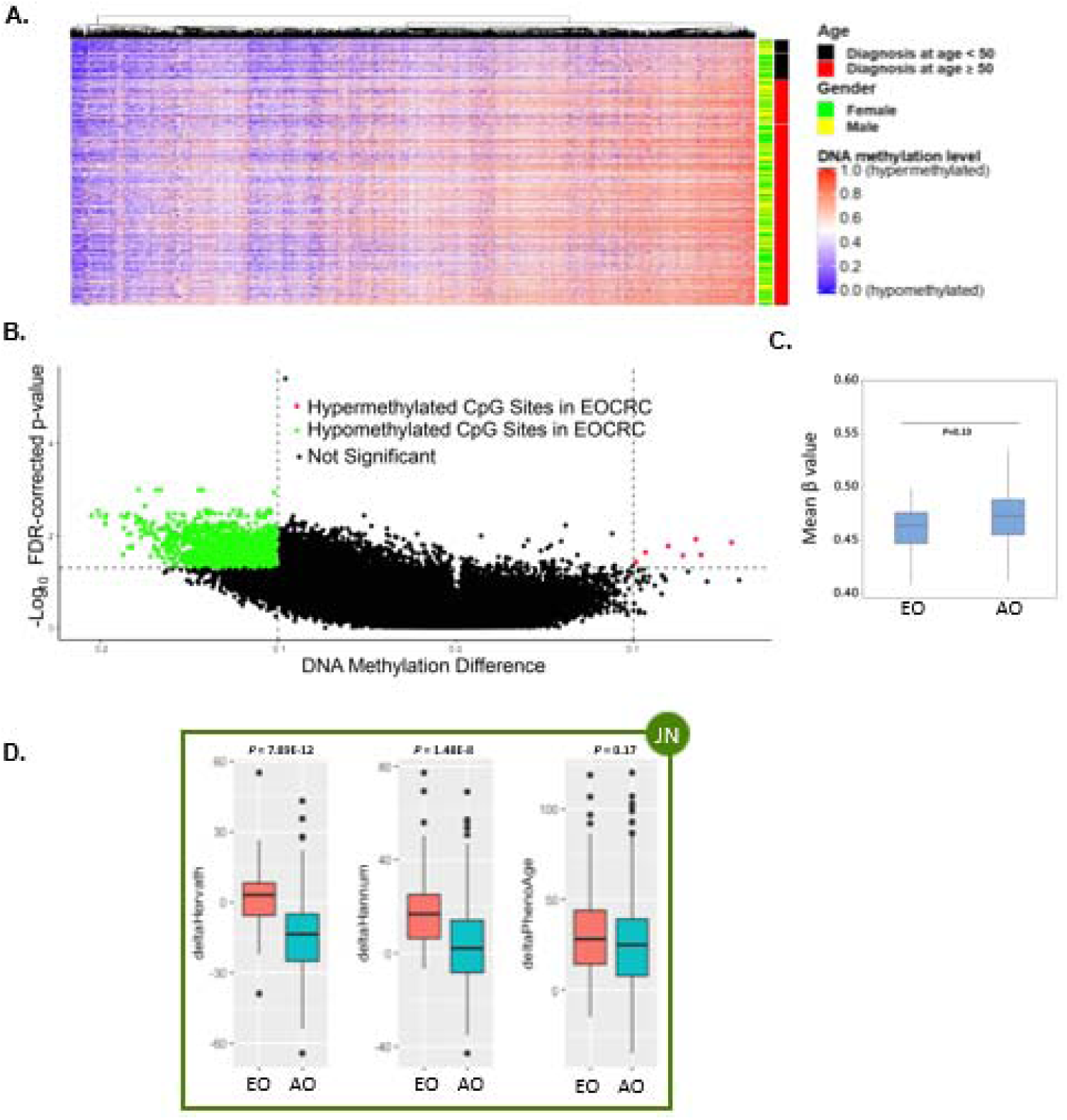
Accelerated aging with epigenetic signatures in EOCRC. **A)** Heatmap was drawn with samples (patients) in the row and CpG-sites (probes) in the column. The DNA methylation values range from 0.0 (completely unmethylated, blue) to 1.0 (completely methylated, red). **B)** The volcano plot demonstrated DMSs (expressed as β values) between EOCRC and AOCRC groups. △β for each CpG site is defined as β value of (EOCRC – AOCRC). False-discovery rate adjusted p < 0.05 and |△β| ≥ 0.1 were used as cut-off values. Sites with △β value ≥ 0.1 and *P* value < 0.05 were defined as significantly hypermethylated sites, which were shown in red; Those with adjusted *P* value < 0.05 and △β value ≤ −0.1, were defined as significantly hypomethylated sites (green). **C)** Box plot showed the mean β value of EOCRC and AOCRC patients was 0.46 ± 0.028 and 0.47 ± 0.027 (*P*-value=0.13, 2-sample *t*-test), respectively. **D)** Boxplots demonstrated the accelerated aging in EOCRC. The DNAm age acceleration (Δ = methylation age – chronological age) in EOCRC and AOCRC for all samples are shown. The Δ_EOCRC_ predicted by Horvath, Hannum, and PhenoAge models were 2.1, 18.4, and 32.8 years. The Δ_AOCRC_ predicted by Horvath, Hannum, and PhenoAge were −14.2, 3.4, and 27.1 years, with *P*-value=7.89E-12, 1.48E-08, and 0.17, respectively.

Prior studies showed that DNA methylation changes through the aging process.^64–66^ To minimize the confounding effect of methylation status solely due to age, we utilized epigenetic clocks to predict the methylation ages between EOCRC and AOCRC patients. Epigenetic clocks are mathematical models to estimate DNA methylation age, which can predict chronological ages. We utilized three different epigenetic models, including Horvath,^61^ Hannum,^62^ and PhenoAge,^63^ to evaluate epigenetic aging in the EOCRC and AOCRC groups. We calculated DNA methylation (DNAm) age acceleration (Δ) as the difference between methylation age and chronological age for each patient (Δ = methylation age – chronological age). The DNAm age acceleration (Δ_EOCRC_) by Horvath, Hannum, and PhenoAge were 2.1, 18.4, and 32.8 years. In comparison, the Δ_AOCRC_ calculated by Horvath, Hannum, and PhenoAge were −14.2, 3.4, and 27.1 years. In other words, both EOCRC and AOCRC groups demonstrated age acceleration (with the exception of AOCRC by Horvath), and the methylation ages from EOCRC were more distant and much older than their chronological ages (*P* value of 7.89 × 10^-12^, 1.48 × 10^-8^, and 0.17, by Horvath, Hannum, and PhenoAge, respectively) **(Figure 1D)**.

### Validation of the Differential Methylation as Affecting Gene Expression

Hypermethylated CpG within the promoter regions commonly leads to the silence of gene transcription and vice versa. To determine whether the DNA methylation differences between EOCRC and AOCRC led to functional effects in the cell (transcription), we integrated differential DNA methylation and gene expression using the starburst plot, which identifies genes whose methylation and expression levels are highly anti-correlated. The starburst plot shows nine distinct quadrants, in which the x-axis shows the FDR adjusted *P*-values for DNA methylation and the y-axis shows the FDR adjusted *P*-values for gene expression. The genes highlighted in the upper left and lower right quadrants show possible gene activation or silencing due to DNA hypomethylation or hypermethylation, with a difference in methylation > 0.1 beta value and a difference in expression with fold change > 2 between EOCRC and AOCRC **(Figure 2A**, per site relationships and *P*-values in **Supplementary Table S2.)**. Among the hypomethylated and up-regulated genes in the EOCRC group, cyclin M1 (CNNM1) and chromogranin A (CHGA) were identified as the key genes whose action may be due to DNA hypomethylation. CNNM1 is associated with neuron cell stemness and self-renewal, and the CHGA gene product modulates the neuroendocrine system.

**Figure 2.**
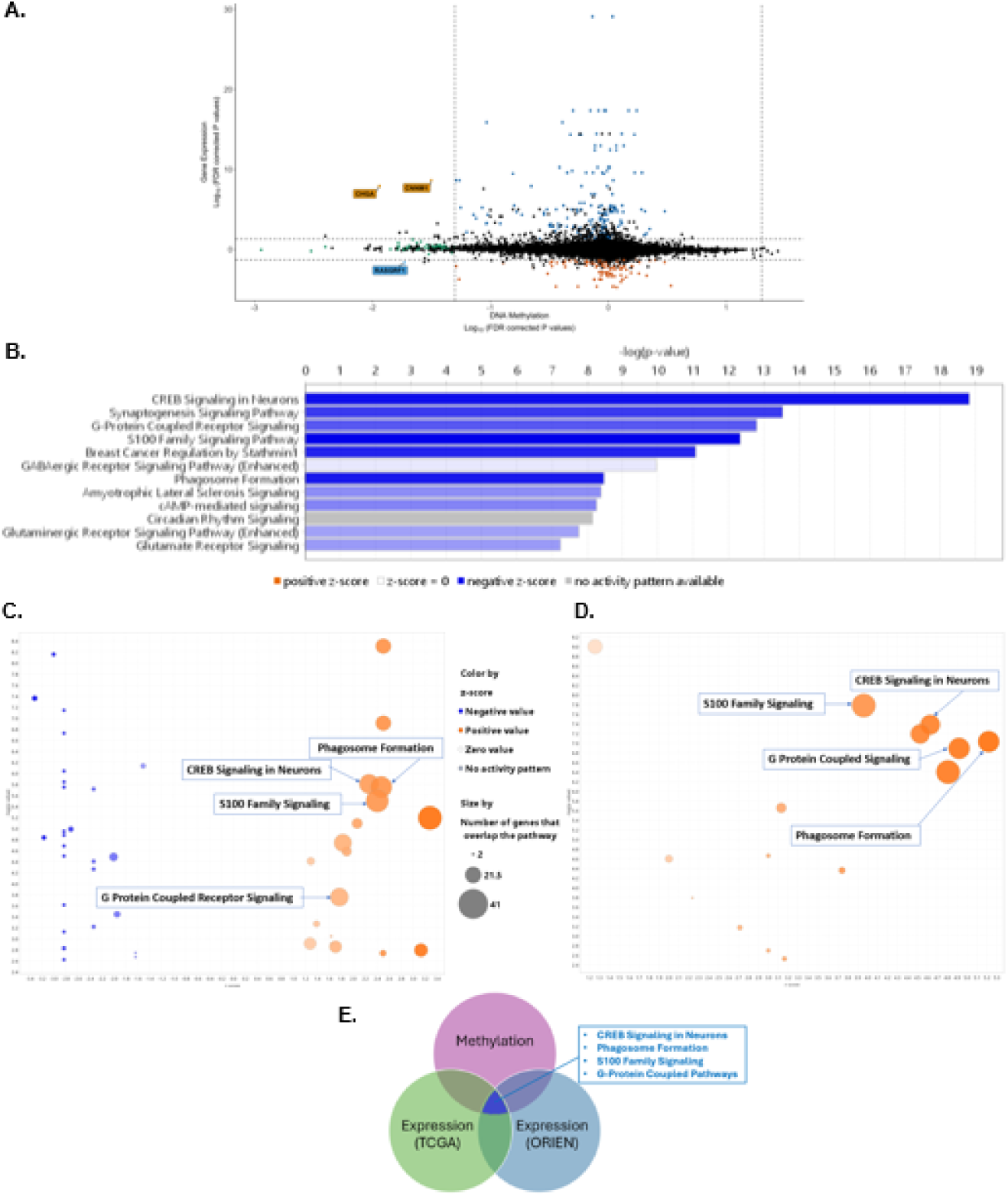
Methylation signature validation through gene expression profiles in EOCRC. **A)** Integration of the differentially methylated sites with paired RNA-seq showed genes that vary in methylation and transcription. The threshold for the FDR-adjusted *P*-values for differences is 0.05 (FDR *P*-value ≤ 0.05). Highlighted genes in orange indicate significant hypomethylation and expression. **B)** The canonical pathways involved with DMGs in EOCRC have an FDR-adjusted *P*-value < 0.05. **C) & D)** Bubble chart of pathways based on DEGs in EOCRC from TCGA and ORIEN, highlighting those predicted by methylation alone. The color of the pathways is based on their z-score, and the size of the bubble is correlated with the number of genes from the dataset that overlap each pathway. **E)** Summary of the analyses to validate the differential methylation, with those supported by all three datasets indicated within the box.

To shed light on accelerated epigenetic aging in EOCRC, it is essential to investigate the biological pathways related to differential methylation in the EOCRC group. A total of 937 hypomethylated genes and 12 hypermethylated genes in EOCRC were identified; we utilized IPA to characterize pathways associated with methylation changes in EOCRC. The top five pathways include cAMP response element binding-protein (CREB) signaling in neurons, synaptogenesis signaling pathways, G-protein-coupled receptor (GPCR) signaling, the S100 family signaling pathway, and breast cancer regulation by Stathmin1 **(Figure 2B)**. These pathways are involved with differentially hypomethylated sites in EOCRC.

Next, we asked whether the pathways involved with differentially methylated genes (DMGs) may modulate cellular transcription. We hypothesized that the pathways identified by the methylation analysis would be important if they also appeared significant from an unsupervised EOCRC vs. AOCRC transcriptional analysis. Four of the pathways found to be important by methylation were found among the most differentially expressed gene (DEG) transcription in the TCGA (**Figure 2C**) and ORIEN (**Figure 2D**) datasets: CREB signaling in neurons, GPCR signaling, phagosome formation, and S100 family signaling pathways, as shown in the Venn diagram (**Figure 2E)**. As a note, these pathways involved with DEGs were shown to be activated in IPA, which suggests that EOCRC-specific methylation patterns may strongly affect cellular transcription.

### Intra-tumoral Microbes Are Not Strongly Associated with EOCRC

Compared to AOCRC, EOCRC patients have unique gene signatures of phagosome formation and S100 family signaling pathways based on canonical pathways associated with DMGs and DEGs. The S100 family proteins are expressed in different cell types (immune and epithelial cells) and regulate cellular processes, including proliferation, inflammation, and invasion.^67–69^ We next sought to determine if the epigenetic aging effects, which appeared to be driven by inflammatory expression programs, could be attributed to another environmentally derived factor: the microbiome. We used the {exotic}^59^ pipeline to count non-human reads in bulk RNA-seq data from TCGA and ORIEN. To validate the presence of the microbes and provide an additional assessment of the EOCRC vs. AOCRC tumor microbe burden, we generated 16S-amplicon data from 60 samples partially overlapping with the ORIEN dataset. The microbes identified in each dataset overlapped, with the most similarity between the RNA-seq datasets (**Figure 3A**). Roughly half of the microbes found in the 16S data were not observed in the RNA-seq results, which may be due to the lower depth expected in bulk RNA-seq data (i.e., greater sensitivity in the 16S data). As an additional validation, we compared the distance between the microbe composition between samples from the same patient (paired, **Figure 3B**) and different patients (not paired). The RNA-seq-derived microbes were more similar to 16S from the same tumor than other CRC samples (Kruskal-Wallis rank sum test *P*-value = 0.02508). Further, the RNA-seq-derived microbes were not significantly different from 16S-derived microbes from tumors or adjacent normal tissue (Kruskal-Wallis rank sum test *P*-value = 0.6682). In the normal samples, the paired samples are more similar than the unpaired samples (*P* = 0.03181). All groups differed significantly from negative controls (all *P*-values < 0.01). This gave us the confidence to analyze the RNA-seq data further for EOCRC and AOCRC differences.

**Figure 3.**
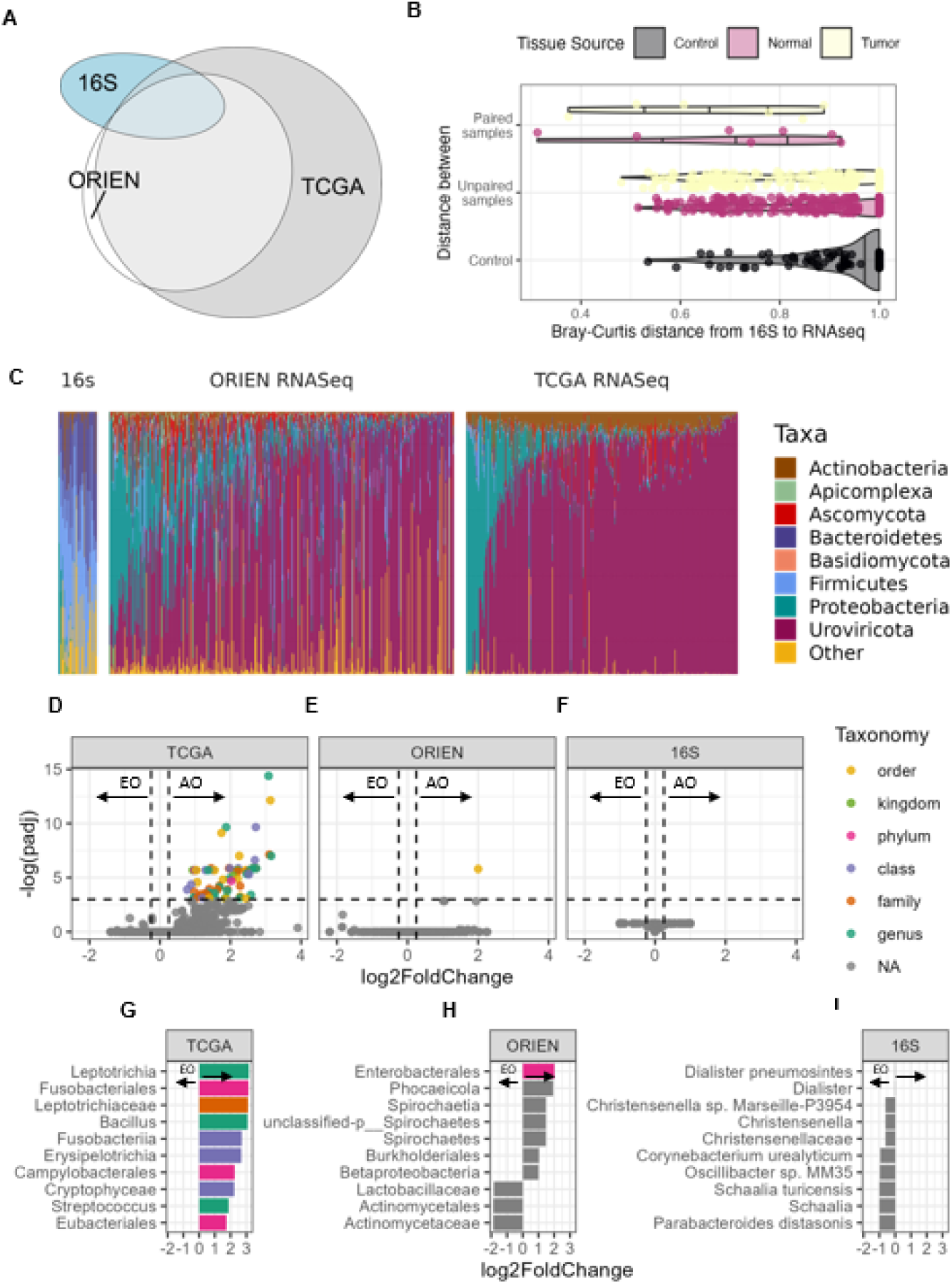
Intra-tumoral microbes do not show consistent differences in EOCRC. **A)** Three datasets (ORIEN, *n* = 353, TCGA, *n* = 357, and 16S, *n* = 30) were analyzed for differences by EOCRC status, which overlapped. **B)** Tumor RNA-seq-derived microbes were most similar to 16S microbes from the same tumor and least similar to adjacent normal tissue from another person. **C)** *Firmicutes* dominated the 16S and RNA-seq microbe abundances, though there were large differences in the number of viruses detected (not detectable by 16S), and the 16S samples showed a more diverse set of phyla (i.e., more “Other”). **D)** In TCGA, microbes were significantly enriched in AOCRC (right), but no microbes passed the significance threshold in EOCRC. **E)** For ORIEN, a single organism was enriched in AOCRC despite a larger sample size. **F)** The 16S data also showed more microbes in the AOCRC samples. **G, H, I)** Effect sizes of the most significant taxa across the three analyses show no similarities in the AOCRC-enriched microbes.

*Firmicutes* dominated the tumor microbes in both the 16S and RNA-seq data. However, there were large differences in the number of viruses detected (not detectable by 16S), and the 16S samples showed a more diverse set of phyla (i.e., more “Other”), likely due to increased sensitivity (**Figure 3C**). Finally, comparing the enrichment of EOCRC vs. AOCRC by abundance showed no microbes enriched in EOCRC across the three datasets (**Figures 3D, E, F,** per microbe log_2_ fold-changes and *P*-values in **Supplementary Tables S3** [TCGA and ORIEN] and **S4** [16S]), including for *Fusobacterium* abundance (**Supplementary Figure S1**). Further, the microbes enriched in AOCRC were different in each dataset (**Figures 3G, H, I**), suggesting no consistent relationship between AOCRC and a particular microbe. Despite a larger sample size, a single organism was enriched in AOCRC for the ORIEN dataset (**Figure 3E, H**). This may be due to the wider geographic distribution from which patient samples were collected relative to the TCGA or 16S datasets.

### Immune-cell Relationship with Microbes in EOCRC

Despite the lack of EOCRC-specific microbe enrichment, the gene signatures in EOCRC led us to explore whether the inflammatory state observed in EOCRC might be due to interactions with microbes. To test this, we estimated all the tumors’ immune cell composition by deconvolution and then correlated the immune cell abundances with microbe abundances at the genus level. Surprisingly, the EOCRC tumors showed stronger correlations than AOCRC in both the ORIEN and TCGA datasets (**Figure 4A**). These correlations were observed across many microbes—rather than just a few strong relationships—consistent with the cohort’s lack of specific microbe enrichment (**Figure 3D-H**). Activated mast cells were the most strongly correlated with many EOCRC microbes consistently between the ORIEN and TCGA datasets (**Figure 4A)**. Specific microbes showed consistent correlations with some immune cells but with a larger scale in EOCRC. For example, *Fusobacterium spp*. was the strongest significantly correlated microbe with neutrophils in both the TCGA and ORIEN datasets and in both EOCRC and AOCRC tumors; however, the size of the EOCRC correlation was larger (**Figure 4B, Supplementary Table S6**). Other associations were inconsistent between the two datasets, with eosinophils and memory B cells most strongly associated with microbes in the ORIEN dataset (**Figure 4A**). At the same time, neutrophils were most strongly correlated in the TCGA dataset (**Figure 4C, Supplementary Table S6**). This suggests that the EOCRC tumors respond to microbes differently and more robustly than AOCRC tumors in a way consistent with innate immune activation and chronic inflammation.

**Figure 4.**
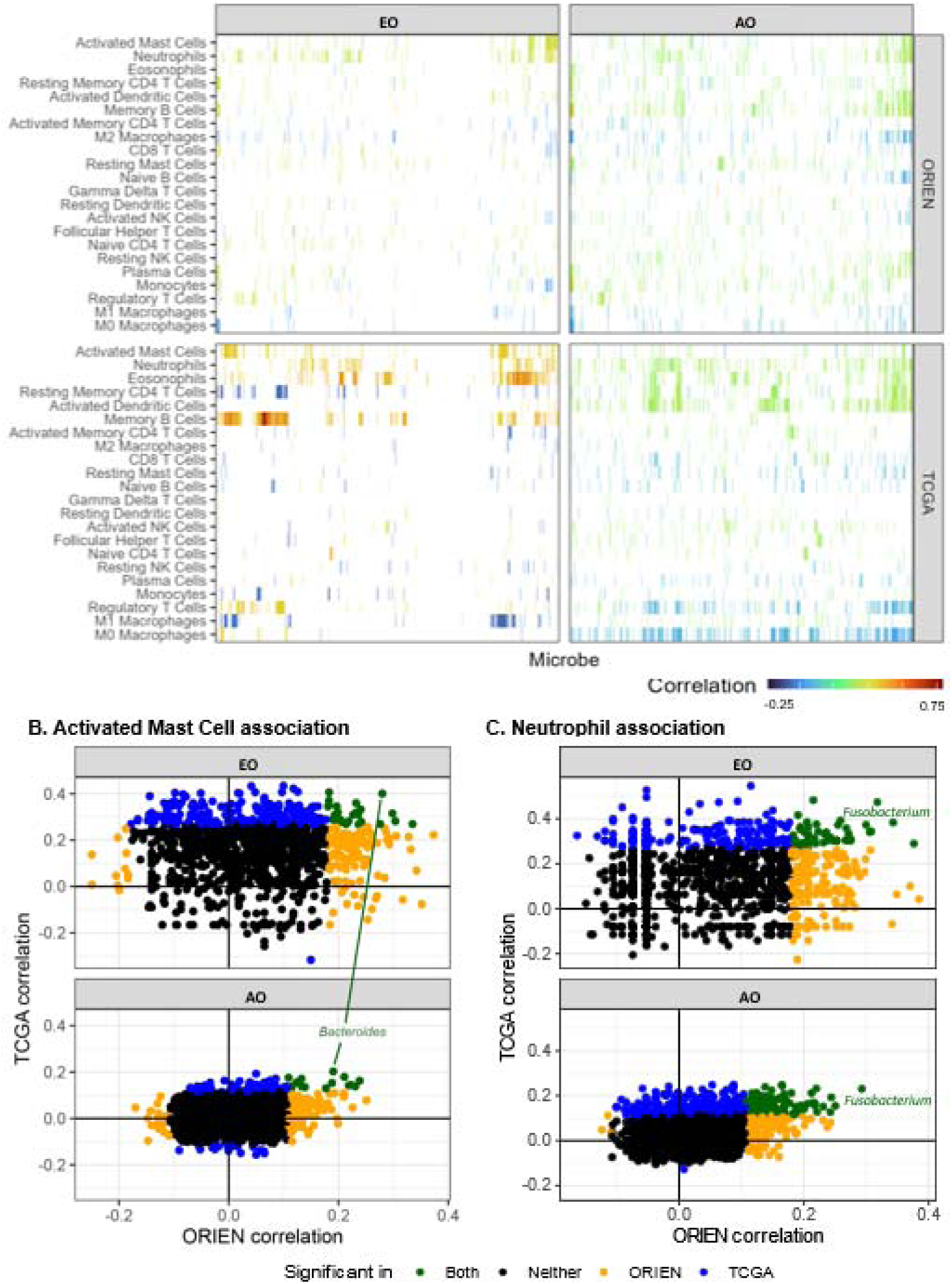
Relationship between microbes and immune cell composition. **A)** All microbe-genus level abundances (columns) correlated to the deconvolved immune-cell abundances (rows), with the strength of the association and its direction indicated by color in the TCGA (top row) and ORIEN (bottom row) datasets. Spearman correlations between the deconvolved abundance of **(B)** activated mast cells or **(C)** neutrophils and microbes in both the TCGA and ORIEN datasets. Concordant associations (significant in both and in the same direction) are shown in green, with select taxa labeled. There were no microbes significantly anti-correlated with activated mast cells or neutrophils.

## DISCUSSION

Epigenetic modifications, including histone modification and DNA methylation, can modulate gene transcription by altering chromatin structure and DNA accessibility without changing the DNA sequence. DNA methylation is not accurately maintained over cell divisions and leads to methylation changes.^70^ Early studies showed that global levels of methylation increase in the infant years and then decrease starting in late adulthood.^71–73^ With the development of microarray and second-generation sequencing technologies, later studies characterized the aging pattern as global methylation loss and focal gains in specific promoter-associated CpG islands.^74–76^ Based on this observation, mathematical models have been developed to utilize these aging-specific CpG sites to predict chronological age and age-related health outcomes, including cardiovascular diseases, neurodegenerative diseases, cancers, and all-cause mortality.

Since sporadic EOCRC is thought to be related to environmental influences and lifestyle changes, we sought to examine methylation alterations in EOCRC. We utilized epigenetic clocks to evaluate if there is a difference in epigenetic aging between EOCRC and AOCRC groups by mining the TCGA Infinium HM450 methylation array database. Our study showed accelerated aging in EOCRC cases compared with AOCRC cases. We found that DNAm age acceleration Δ_EOCRC_ (methylation age – chronological age) computed by different epigenetic clock models was 12 years older than Δ_AOCRC_patients. Certainly, there are variations in predicted methylation ages using different epigenetic models, and it will remain challenging to select the best model to evaluate accelerated aging in EOCRC.

Our result is consistent with the study findings from Buchanan’s group, in which they demonstrated accelerated aging in EOCRC using their unique methylation signature comprising 234 methylation loci.^77^ Our analysis demonstrated that the majority of DMGs are hypomethylated with the unique methylation signatures in EOCRC cases, including CREB signaling in neurons, synaptogenesis signaling pathways, GPCR signaling, phagosome formation, S100 family signaling pathway, and breast cancer regulation by Stathmin1. Correspondingly, canonical pathways involved with differential GEs are partially overlapping with the DNA methylation profile in EOCRC, including CREB signaling in neurons, GPCR signaling, phagosome formation, and S100 family signaling pathway. Pathways involved with DEGs were upregulated, corresponding to hypomethylated DMGs. These overlapping pathways between DNA methylation and gene expression indicate that epigenetic modulation (influenced by environmental risk factors) may lead to the development of EOCRC.

It is well known that epigenetic alterations are one of the hallmarks of aging,^78^ and neurodegenerative diseases may cause accelerated aging by inducing DNA damage, oxidative stress, inflammation, and metabolic dysregulation.^79–82^ CREB, a transcription factor, acts as a downstream effector of the GPCR-protein kinase A (PKA)-CREB signaling pathway. CREB has been shown to play an essential role in promoting cell proliferation, neuronal survival, and synaptic plasticity in the central nervous system,^83–85^ and dysregulated CREB signaling may be associated with cognitive deficits and aging.^86^ GPCR/PKA/CREB signaling has been shown to play a role in maintaining stemness, promoting progression and metastasis, and CREB knockdown reduces the metastatic potential in CRC.^87, 88^ However, it is unknown whether DNA methylation plays a role in regulating GPCR/PKA/CREB signaling in EOCRC and accelerated aging.

The S100 family proteins are involved with the cell cycle, cytoskeleton activity, cell differentiation and motility, inflammation, and/or antimicrobial activity.^89^ They are associated with neurological disorders, cancers, and cardiac and inflammatory diseases.^90^ Several S100 proteins, including S100A1, S100A2, S100A4 and S100A11, were found in colon carcinomas with more aggressive disease and worse clinical outcomes.^91, 92^ S100A4 can activate NF-κB signaling and the Wnt/β-catenin pathway to promote tumorigenesis and metastasis in CRC.^93, 94^ Knockdown of the S100 protein by lentiviral RNA interference can inhibit colon cancer growth and metastasis.^95^ Earlier studies also demonstrated that DNA methylation modulated gene transcription regulation in S100 functions in different cancers, including colon cancer.^96, 97^ Supported by these abundant research findings, we plan to investigate the S100 family signaling pathway in EOCRC in future studies.

Gut microbiota play a central role in various host physiological and metabolic functions, including fiber degradation and fermentation, energy supply, lipid metabolism, vitamin synthesis, and maintenance of intestinal barrier integrity.^98–101^ Intestinal dysbiosis can cause an increased inflammation state, produce toxic metabolites, and promote an immunosuppressive tumor microenvironment that suppresses antitumor immune surveillance.^102–106^ Chronic inflammation activates key pro-proliferative signaling pathways via NF-κB, resulting in aberrant proliferation and epithelial cell transformation, thereby promoting carcinogenesis.^40, 41^ It has been established that certain pathogenic microbes, including *Escherichia coli*, *Enterococcus faecalis*, *Bacteroides fragilis* and *Fusobacterium nucleatum* are increased in CRC patients.^107^ Therefore, the link between a Western diet and dysbiosis has stimulated considerable interest in uncovering alterations in microbial community structure in EOCRC. Several recent studies, however, failed to identify enrichment of *Fusobacterium nucleatum* in the EOCRC patient population, which is consistent with our study.^108–110^ Our results did not identify enrichment of specific microbes in EOCRC across both the TCGA and ORIEN datasets. Further studies are required to determine whether intestinal dysbiosis may have a direct association with EOCRC.

Interestingly, when comparing the microbes to inferred immune cell abundances, the EOCRC microbes showed more, larger correlations with immune cells despite the same immune cell-microbe relationships being found in EOCRC and AOCRC (e.g., *Fusobacterium* was consistently associated with neutrophils in both EOCRC and AOCRC). This implies that EOCRC may respond more strongly to the presence of microbes, driving a pro-inflammatory phenotype, inflammaging, and earlier cancer development. Further, this suggests an approach to increase immune tolerance to gut microbes that may protect against EOCRC.

Our study has several limitations. We utilized the TCGA and ORIEN datasets for methylation and microbiome analysis to use a discovery and validation approach. However, technical differences between the datasets may have led to a higher false negative rate than a single, larger dataset with uniform data generation procedures and processing. Another limitation is the lack of microbiome source material, likely from the gut and responding to diet and lifestyle factors. A thorough treatment would include paired collection of stool as well as mucosa-adherent and luminal microbiome samples. In addition, a set of normal colonic tissue samples from healthy donors or paired normal adjacent tissue would be ideal to serve as controls to evaluate cancer-associated microbes and epigenetic aging. Further, the study remains strictly correlative. Our findings need further validation in large cohorts of patients, as well as mechanistic studies in animal models to identify causality.

## CONCLUSIONS

We found accelerated aging in EOCRC by mining the TCGA datasets. Methylation ages predicted by three different epigenetic clocks in EOCRC patients were 12 years older on average than methylation ages in AOCRC patients. Canonical pathways involved with differential methylation and gene expression partially overlapped, including CREB signaling in neurons, GPCR, phagosome formation, and S100 family signaling pathways in EOCRC. These four pathways were found to be hypomethylated and activated in gene transcription in both TCGA and ORIEN datasets. There was no significant intra-tumoral microbiome identified in EOCRC. However, specific microbes (such as *Fusobacterium*) showed consistent correlations with certain immune cells (neutrophiles) with a larger scale in EOCRC. As a summary, our findings suggested that DNA methylation changes and epigenetic aging may contribute to the development of EOCRC.

## ACKNOWLEDGMENTS

This project was partly supported by the Paul Calabresi Career Development Award for Clinical Oncology (K12 Research Training K12CA133250) (Ning Jin); The Ohio State University Comprehensive Cancer Center and the NIH (P30CA016058); The Ohio State University Center for Clinical and Translational Science and the National Center for Advancing Translational Sciences (8UL1TR000090-05); The Huntsman Cancer Institute, Comprehensive Cancer Center at the University of Utah and the NIH (P30CA042014 & U01 CA206110 [Ulrich, Hardikar, Figueiredo]); an Alpha Omega Alpha Carolyn L. Kuckein Student Research Fellowship (D.J. Grencewicz); a Samuel J. Roessler Memorial Scholarship (D.J. Grencewicz); the National Institute on Aging (K01AG070310 to DS); the American Cancer Society (to DS). This work was also supported by the Pelotonia Institute of Immuno-Oncology. The authors acknowledge the support and resources of the Ohio Supercomputing Center (PAS1695 and PCON0005). The results published here are in whole or part based upon data generated by TCGA Research Network: https://www.cancer.gov/tcga. We would like to thank Angela Dahlberg, Editor, Division of Medical Oncology at The Ohio State University Comprehensive Cancer Center, for editing and proofreading the manuscript.

## Conflicts of Interest

**Eric A Singer**: Astellas/Medivation: research support (clinical trial); Johnson & Johnson: advisory board; Merck: advisory board; Vyriad: advisory board; Aura Biosciences: data safety monitoring board

**John D Carpten**: Roche/Genentech

**Carlos HF Chan**: None related to this project. Other unrelated projects and clinical trials: Research support from Checkmate Pharmaceuticals, Regeneron, Angiodynamics, Optimum Therapeutics

**Gregory Riedlinger**: AstraZeneca advisory board

**Bryan P Schneider**: Genentech-Research support (drug supply only); Pfizer-Research support (Drug supply only); Foundation Medicine-research support (sequencing support)

**Ahmad A Tarhini**: Contracted research grants with institution from Bristol Myers Squib, Genentech-Roche, Regeneron, Sanofi-Genzyme, Nektar, Clinigen, Merck, Acrotech, Pfizer, Checkmate, OncoSec. Personal consultant/advisory board fees from Bristol Myers Squibb, Merck, Easai, Instil Bio Clinigin, Regeneron, Sanofi-Genzyme, Novartis, Partner Therapeutics, Genentech/Roche, BioNTech, Concert AI, AstraZeneca outside the submitted work.

**Yousef Zakharia**: Advisory Board: Bristol Myers Squibb, Amgen, Roche Diagnostics, Novartis, Janssen, Eisai, Exelixis, Castle Bioscience, Genzyme Corporation, Astrazeneca, Array, Bayer, Pfizer, Clovis, EMD Serono, Myovant; Grant/research support from: Institution clinical trial support from NewLink Genetics, Pfizer, Exelixis, Eisai; DSMC: Janssen Research and Development; Consultant honorarium: Pfizer, Novartis

**Cornelia M Ulrich**: As cancer center director, Cornelia Ulrich oversees research funded by several pharmaceutical companies but has not received funding directly herself.

The remaining authors declare no conflict of interest.

## Author Contributions

Conceptualization: NJ Resources: all authors

Data curation: Ulrich CM, RH, CD,

Software: CD, RH, MJS, YL,

Formal analysis: CD, RH, MJS, YL, SY,

NJ Supervision: NJ, DS

Funding acquisition: Ulrich CM

Validation: SY

Investigation: CD, RH, MJS, YL, SY

Visualization: RH, CD, SY

Methodology: DS, NJ

Writing - original draft: NJ

Project administration: NJ

Writing - review and editing: all authors

