## Supplementary figures and images for "Epigenetic Modulation, Intra-tumoral Microbiome and Immunity in Early Onset Colorectal Cancer"

### Figure S1

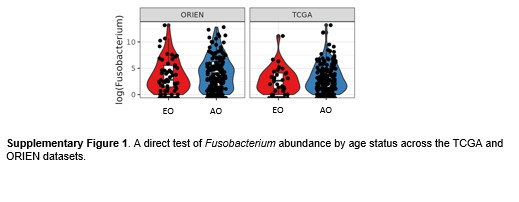
